# Asynchronous microexon splicing of *LSD1* and *PHF21A* during neurodevelopment

**DOI:** 10.1101/2024.03.21.586181

**Authors:** Masayoshi Nagai, Robert S. Porter, Elizabeth Hughes, Thomas L. Saunders, Shigeki Iwase

## Abstract

LSD1 histone H3K4 demethylase and its binding partner PHF21A, a reader protein for unmethylated H3K4, both undergo neuron-specific microexon splicing. The LSD1 neuronal microexon weakens H3K4 demethylation activity and can alter the substrate specificity to H3K9 or H4K20. Meanwhile, the PHF21A neuronal microexon interferes with nucleosome binding. However, the temporal expression patterns of LSD1 and PHF21A splicing isoforms during brain development remain unknown. In this work, we report that neuronal PHF21A isoform expression precedes neuronal LSD1 isoform expression during human neuron differentiation and mouse brain development. The asynchronous splicing events resulted in stepwise deactivation of the LSD1-PHF21A complex in reversing H3K4 methylation. We further show that the enzymatically inactive LSD1-PHF21A complex interacts with neuron-specific binding partners, including MYT1-family transcription factors and post-transcriptional mRNA processing proteins such as VIRMA. The interaction with the neuron-specific components, however, did not require the PHF21A microexon, indicating that the neuronal proteomic milieu, rather than the microexon-encoded PHF21A segment, is responsible for neuron-specific complex formation. These results indicate that the PHF21A microexon is dispensable for neuron-specific protein-protein interactions, yet the enzymatically inactive LSD1-PHF21A complex might have unique gene-regulatory roles in neurons.

## Introduction

Brain development, the process in which the organ governing cognition forms, is arguably one of the most complex and intricate developmental processes. Precisely regulated gene expression, including transcriptional regulation, is key for normal brain development. Human genetics studies of neurodevelopmental disorders have highlighted the critical roles of chromatin regulation in brain development. Chromatin regulators, which place, read, and erase histone and DNA modifications, represent a major gene group responsible for neurodevelopmental disorders, such as intellectual disability, schizophrenia, and autism spectrum disorders (1–3). However, chromatin regulators are broadly expressed across the body, making it challenging to understand the brain’s particular vulnerability to chromatin dysregulation.

Recent work has begun to unveil the unique features of chromatin regulations in neurons. For example, non-CpG DNA methylation, primarily in the CA context, is found abundantly in neurons undergoing synaptogenesis and modulates synaptic gene expression (4, 5). Our group has reported evolutionally conserved neuron-specific microexon splicing events in 14 chromatin regulators (6). In most cases, except for the LSD1 and PHF21A, as discussed below, the functional consequences of neuronal splicing events remain unknown. Thus, these observations have opened an area of research to investigate the roles and mechanisms of neuron-specific chromatin regulation and its potential link to human neurodevelopmental disorders.

Of the 14 chromatin factors that undergo neuron-specific splicing, two proteins—histone H3K4 demethylase LSD1 (aka KDM1A) and H3K4me0 reader protein PHF21A (aka BHC80)—are particularly intriguing for two reasons. First, their non-neuronal forms assemble into a stoichiometric complex suppressing neuron-specific genes in non-neuronal cells (7, 8). The complex consists of LSD1, PHF21A, class I histone deacetylases HDAC1 and HDAC2, HMG-box DNA binding protein BRAF35, and CoREST, which promote the nucleosome binding of LSD1 (7–11). REST/NRSF is the transcription factor that recruits the PHF21A-LSD1 complex to neuron-specific genes in non-neuronal cells for transcriptional silencing (7). Second, loss-of-function mutations in both genes, *KDM1A* and *PHF21A*, cause rare neurodevelopmental disorders that involve cognitive deficits (12–14). Since these mutations impact both canonical and neuronal isoforms of LSD1 and PHF21A, it remains to be determined which isoform is responsible for observed cognitive deficits.

The neuronal microexon splicing events in LSD1 and PHF21A cause small—just a few encoded amino acids—yet significant changes in their protein functions. The LSD1 neuronal splicing reduces H3K4 demethylation activity (15) and possibly alters substrate specificity towards H3K9 (16) or H4K20 (17). Furthermore, the LSD1-Thr369, one of the four amino acids encoded by the neuronal microexon, is phosphorylated in the brain and, in turn, negatively modulates binding to CoREST and HDAC1/2 (18). The LSD1 neuronal splicing is modulated by neuronal activity (19), thereby contributing to excitation-inhibition balance, stress-response behavior (20), and learning and memory (17). Meanwhile, PHF21A neuronal splicing ablates the DNA binding function of PHF21A mediated by an AT-hook motif, which is present in the canonical (PHF21A-c) but not neuronal PHF21A isoform (PHF21A-n) (6). In sum, the emerging understanding is that the neuronal splicing events in LSD1 and PHF21A interfere with H3K4 demethylating function and contribute to the circuit homeostasis and cognitive functions.

Critical questions regarding LSD1 and PHF21A neuronal splicing still need to be addressed. First, developmental expression patterns of LSD1 and PHF21A isoforms have yet to be examined in detail. Second, we do not know whether neuronal LSD1 and PHF21A isoforms participate in a similar complex as non-neuronal cells or whether the neuronal complex is unique. To address these questions, in this study, we characterized the expression kinetics of LSD1 and PHF21A neuronal isoforms during neuronal differentiation and its consequences on the demethylase activity of the complex. We employed a proteomics approach to test whether LSD1 and PHF21A assemble a neuron-specific complex and further provided insights into the functional roles of neuronal splicing in the complex assembly by generating a mouse mutant in which PHF21A-n is replaced with PHF21A-c in neurons.

## Results

### PHF21A completes switching to neuronal form prior to LSD1

To examine the expression kinetics of PHF21A-n and LSD1-n during neuronal differentiation, we first turned to *in vitro* differentiation of Lund human mesencephalic (LUHMES) cells into neurons. LUHMES cells are known to produce a homogeneous neuronal population quickly once their differentiation is induced by dibutyryl-cAMP, thereby making them well-suited for biochemical investigations (21). The reverse transcriptase coupled polymerase chain reaction (RT-PCR) indicated that *PHF21A* mRNA completely switched from *PHF21A-c* to *PHF21A-n* after 3 days of neuronal differentiation (Fig. 1A). In contrast, the appearance of *LSD1-n* form began only after 6 days of differentiation. *LSD1-c* continues to be expressed during neuronal differentiation, which is consistent with previous reports (15).

**Figure 1:**
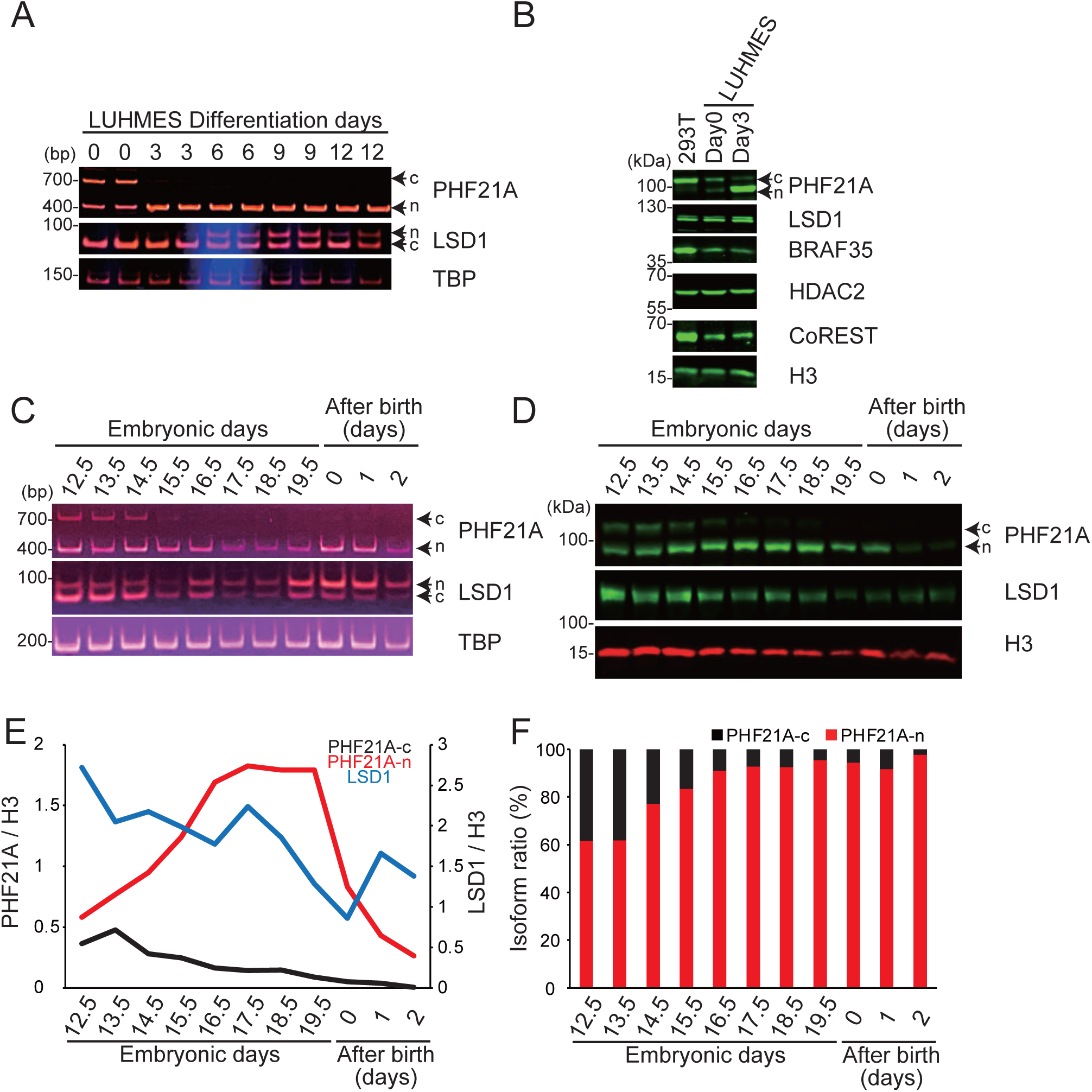
The expression of PHF21A-n increases faster than that of LSD1-n. **(A)** mRNA levels of LSD1 and PHF21A isoforms in LUHMES cells. Cells were differentiated into neurons as indicated and harvested from day 3 to day 12, analyzed by RT-PCR (n=2). **(B)** Expression of PHF21A and associated proteins in LUHMES cells and 293T cells examined by Western blot analysis using antibodies as indicated. **(C)** mRNA levels of LSD1 and PHF21A isoforms in the developing mouse brain. Whole-cell lysates prepared from mouse whole brains (E12.5 and E13.5) and cortices (from E14.5 to P2) at indicated periods were analyzed by RT-PCR. **(D)** Expression of PHF21A and LSD1 proteins in the developing mouse brain. Whole-cell lysates were subjected to Western blot analysis. **(E)** Quantification of Western signals for PHF21A-c, PHF21A-n, and LSD1 normalized by histone H3. **(F)** The ratio of PHF21A protein isoforms in the developing mouse brain based on the Western signals.

To measure the *LSD1*-*n/c* ratio more quantitatively, we performed Illumina-based complete amplicon sequencing of the *LSD1* RT-PCR products harboring the alternative sequences internally. We adjusted the isoform ratio by accounting for the PCR efficiency difference; *LSD1-c* efficiency was 2.8% greater than *LSD1-n* PCR efficiency, which translates to a 1.62-fold underestimation of *LSD1-n* abundance in amplicon sequencing carried out with a 30cycle PCR (Fig. S1A & S1B, and see Material & Method). The LSD1-c:LSD1-n ratio obtained by complete amplicon sequence described above was then adjusted with the correction values. *LSD1-n* emerged on day 3, kept increasing, and reached a plateau at approximately 37.7% on day 9 of LUHMES cell differentiations (Fig. S1C). A western blot using an anti-PHF21A antibody showed that the PHF21A-c protein was mostly replaced with PHF21A-n on day 3 (Fig. 1B), which agreed well with the RT-PCR result. Note that WTN cannot distinguish LSD1-c and LSD1-n, which only differ in 4 amino acids; the doublet signal in WTN originates from another alternative sequence located at the region close to the LSD1 N terminus (22). The large difference in *PHF21-n* and *PHF21A-c* PCR efficiency (23%) precluded a reasonably accurate quantification of the *PHF21A* isoform ratio. These data indicated that the expression of PHF21A-n precedes that of LSD1-n in differentiating LUHMES cells.

We next examined the expression kinetics of the PHF21A and LSD1 isoforms in developing mouse brains. RT-PCR indicated that *PHF21A* mRNA completely switched from *PHF21A-c* to *PHF21A-n* after E15.5 mouse cortex (Fig. 1C). In contrast, the LSD1-c was present throughout the developmental periods examined. In this RT-PCR analysis, the LSD1-n/c ratio appeared to show step-wise increases on E16.5 and P0. Quantification of the results with Ilummina complete amplicon sequence indicated that *LSD1-n* mRNA consists of 42% at E18.5 and increased further to 81% at P0 brain (Fig. S1D). In contrast, similar to *PHF21A* mRNA isoforms, the percentage of PHF21A-n protein reached a plateau (91%) already in the E16.5 mouse cortex (Fig. 1D, E, & F). Notably, total protein levels of LSD1 and PHF21A decreased over the examined time periods, implicating a greater contribution of these factors to embryonic brain development than brain maturation after birth. These results indicate that similar to the observation in LUHMES cells, the increase of neuronal form composition is faster for PHF21A than LSD1 *in vivo*. Since PHF21A adopts the neuronal form earlier than LSD1, the two proteins potentially form an intermediate complex containing PHF21A-n and LSD1-c prior to the mature neuronal complex containing PHF21A-n and LSD1-n, especially in the late gestation period.

### The intermediate complex with PHF21A-n and LSD1-c demethylates H3K4

Prior studies have described LSD1-n as a weaker H3K4 demethylase (15), H3K9-(16), and H4K20-(17) demethylase. Having established the differential expression kinetics of PHF21A and LSD1 isoforms, we sought to determine the substrate specificity of the intermediate complex containing PHF21A-n and LSD1-c. We immunoprecipitated (IP) the complexes with a PHF21A antibody from undifferentiated (day 0) and differentiating (day 3) LUHMES cells. In undifferentiated LUHMES cells on day 0, 62.8% of PHF21A and 100% of LSD1 adopt the canonical form (Fig. 1B and Fig. S1C). On day 3, 91.6% of PHF21A adopts the neuronal form (Fig. 1B), while 94% of LSD1 is still the canonical form (Fig. S1C). PHF21A-IP samples from both day 0 and day 3 contained LSD1, indicating that, like the PHF21A-c, PHF21A-n interacts with LSD1-c, forming the predicted intermediate complex.

We then incubated the IP samples with recombinant designer mono-nucleosomes carrying either H3K4me2, H3K9me2, or H4K20me2. Demethylation reactions can be detected by either the decrease of di-methylation or the appearance of mono-methylation after the reaction via Western blot analysis, as previously described (Fig. 2A) (6). As a control, we examined the enzymatic activity of PHF21A-complex from 293T cells, which only express PHF21A-c and LSD1-c. We reliably detected the generation of H3K4me1 by the PHF21A complexes isolated from both day 0 and day 3 LUHMES cells as well as 293T cells (Fig. 2B). The H3K4me1 signals were absent or nearly absent when the H3K4me2-nucleosome was omitted from the reaction, ruling out the contribution of co-precipitated cellular H3K4me1-nucleosome to WTN signals (Fig. 2B, lanes 6, 10, &14). The control IgG IP samples did not show any activity (Fig. 2B, lanes 4, 8, and 12). In this LUHMES cell system, the overall abundance of PHF21A was greater during differentiation compared to the undifferentiated state, leading to a greater LSD1 level in IP samples, although the total LSD1 level was unchanged during differentiation (Fig. 1C). With the greater LSD1 level, the day 3-complex yielded a similar level of H3K4me1 compared to the day 0-complex, suggesting weaker enzymatic activity of the intermediate complex with PHF21A-n and LSD1-c compared to the canonical complex with PHF21A-c and LSD1-c.

**Figure 2:**
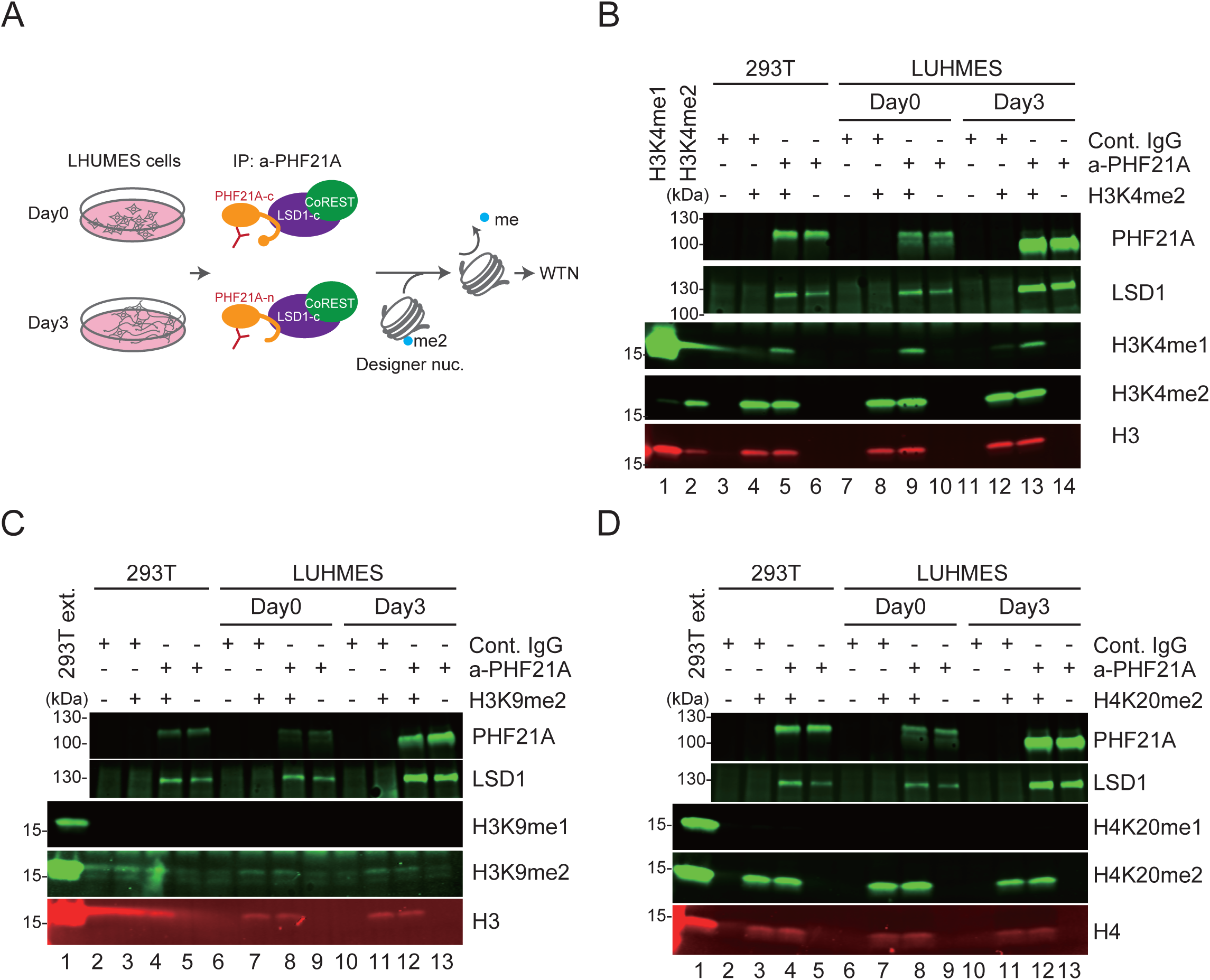
A weaker H3K4 demethylation activity of the intermediate complex with PHF21A-n and LSD1-c. **(A)** Schematic of the demethylation assay. The PHF21A-containing complex was immunoprecipitated from LHUMES cell nuclear extracts and incubated with designer nucleosomes with specific lysine di-methylations. **(B-D)** Western blot using the antibodies for indicated histone methylations to detect demethylase activity. The reactions were carried out using H3K4me2-**(B)**, H3K9me2-**(C)**, and H4K20me2-**(D)** nucleosomes. The appearance of mono-methylated lysine indicates the demethylation activity. The abundance of LSD1, PHF21A, and total H3 (c-term) were examined with specific antibodies. H3K4me1/me2 designer nucleosomes and 293T nuclear extract serve as specificity and positive control for the histone antibodies, respectively.

With H3K9me2 or H4K20me2 nucleosomes, we did not find either the decrease of the di-methylation signal or the appearance of mono-methylation, indicating that the complexes were inactive to these substrates (Fig. 2C and 2D). These results demonstrate that the substrate of the developmental intermediate complex with PHF21A-n and LSD1-c is H3K4.

### No detectable demethylation activity of the mature neuronal complex with PHF21A-n and LSD1-n

In our previous work with reconstituted LSD1-CoREST-PHF21A tripartite complex, we found that the mature complex with LSD1-n and PHF21A-n has weaker H3K4 demethylase activity and lacks detectable activity to H3K9me2 and H4K20me2 (6). However, the reconstitution experiment with purified proteins left a possibility that additional interaction partners could bestow new substrate specificity to the neuronal complex. Indeed, a previous report showed that the interaction with SVIL protein in neurons led LSD1-n to demethylate H3K9 instead of H3K4 (16). We, therefore, sought to determine the substrate specificity of the mature complex with PHF21A-n and LSD1-n isolated from neurons.

Our initial attempt was to differentiate LHUMES cells further than day 3 to the point where a significant fraction of LSD1 adopts neuronal form. However, the LSD1-n level did not reach >40% and reached a plateau on day 9, and subsequent culture did not result in a greater LSD1-n ratio, as discussed earlier (Fig. S1C). We also noted that prolonged culture of LUHMES cells led to fewer cell numbers and made it impractical to collect sufficient IP materials for demethylation assays. For these reasons, we turned to the mouse P0 brains, which show high expression levels of the two proteins adopting predominantly neuronal forms; 94% of PHF21A and 81% of LSD1 are neuronal forms (Fig. 1F and Fig. S1D).

We carried out PHF21A IP from the P0 mouse cortices and demethylation assay using the recombinant designer nucleosomes (Fig. 3A-C). Unlike the intermediate complex in Fig. 2, we did not detect the specific appearance of H3K4me1 when both the H3K4me2 nucleosomes and the complex were present in the reaction (Fig. 3A, compare lane 9 to lanes 8 & 10). Meanwhile, the PHF21A complex from 293T cells shows the specific H3K4me1 appearance (Fig. 3A, lane 5) in the parallel reactions. Importantly, LSD1 levels in PHF21A-IP samples were greater in the brain samples compared to 293T, ruling out the possibility that an insufficient amount of LSD1-PHF21A complex resulted in the lack of enzymatic activity. When we used the H3K9me2 and H4K20me2 nucleosomes, the P0 brain PHF21A complex did not consistently generate mono-methylations either (Fig. 3B and 3C). These results indicate that the mature neuronal complex with LSD1-n and PHF21A-n isolated from the brain lacks detectable enzymatic activity in these assays.

**Figure 3:**
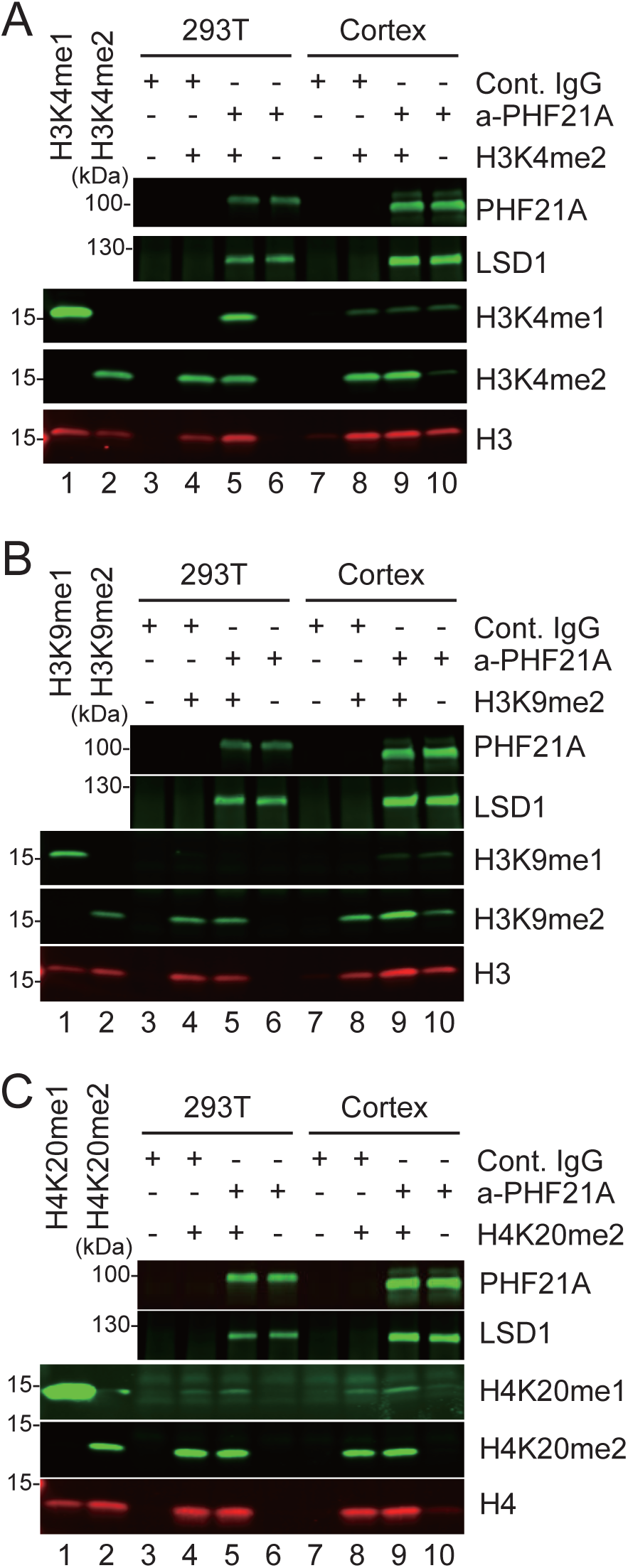
No detectable demethylation activity of the mature neuronal complex with PHF21A-n and LSD1-n. The demethylation assay of immunoprecipitated PHF21A complexes from the P0 mouse cortices using the designer nucleosomes carrying H3K4me2 **(A)**, H3K9me2 **(B)**, and H4K20me2 **(C**). Neither a reduction of di-methylation nor the appearance of mono-methylation was observed with any demethylation reactions.

### Comparative proteomics analysis of PHF21A-containing complex in MEFs and neurons

LSD1-n interacts strongly with SVIL protein when SH-SY5Y cells are differentiated into neurons (16). SVIL is the only protein that exhibits higher binding to LSD1 in differentiated SH-5SY cells compared to undifferentiated SH-SY5Y cells (16). However, the splicing pattern of PHF21A is unknown in SH-SY5Y cells. We reason that investigating PHF21A-n-interacting proteins in neurons might provide insights into the undetectable demethylation activity of the mature complex.

To this end, we performed a comparative PHF21A immunoprecipitation coupled with mass spectrometry (IP-MS) study using mouse embryonic fibroblast (MEF) and cortical neuron cultures (DIV7) (Fig. 4A). LSD1-n is 91% of total LSD1 in the cortical neurons, while absent in MEF (Fig. S1E). PHF21A is predominantly the canonical form in MEF and neuronal form in cortical neurons (Fig. 4F). Though proteins specifically precipitated by the PHF21A antibody were barely visible in silver staining (Fig. 4A), we were able to identify consistently co-precipitated proteins across replicates (n = 3) (Table S1). Of the total 108 proteins identified, 8 proteins in MEF and 17 proteins in cortical neurons passed our threshold: p< 0.1, FC (anti-PHF21A/control IgG) > 1.5, and peptide number ≥ 6 (Fig. 4B and 4C). Notably, all 8 proteins identified in MEF were also identified in neurons (Fig. 4D). Most of these common interactants, such as LSD1, RCOR1 (aka CoREST), HDAC2, and HMG20B (aka BRAF35), were known components of previously isolated LSD1-complexes (7, 8, 23, 24). In neurons, 9 additional proteins were identified. Of these, zinc-finger transcription factors MYT1 and MYT1L are exclusively expressed in embryonic neurons (25) (Fig. 4E). Primary roles of some neuron-specific PHF21A-interaction partners are outside chromatin regulation; DDX5 and VIRMA have been implicated in splicing and mRNA methylation (26–31), while TUBB5, TUBA1A, ACTA2, and MYH10 comprise of the cytoskeleton (32–35), which requires further validation of interaction (See Fig. 5).

**Figure 4:**
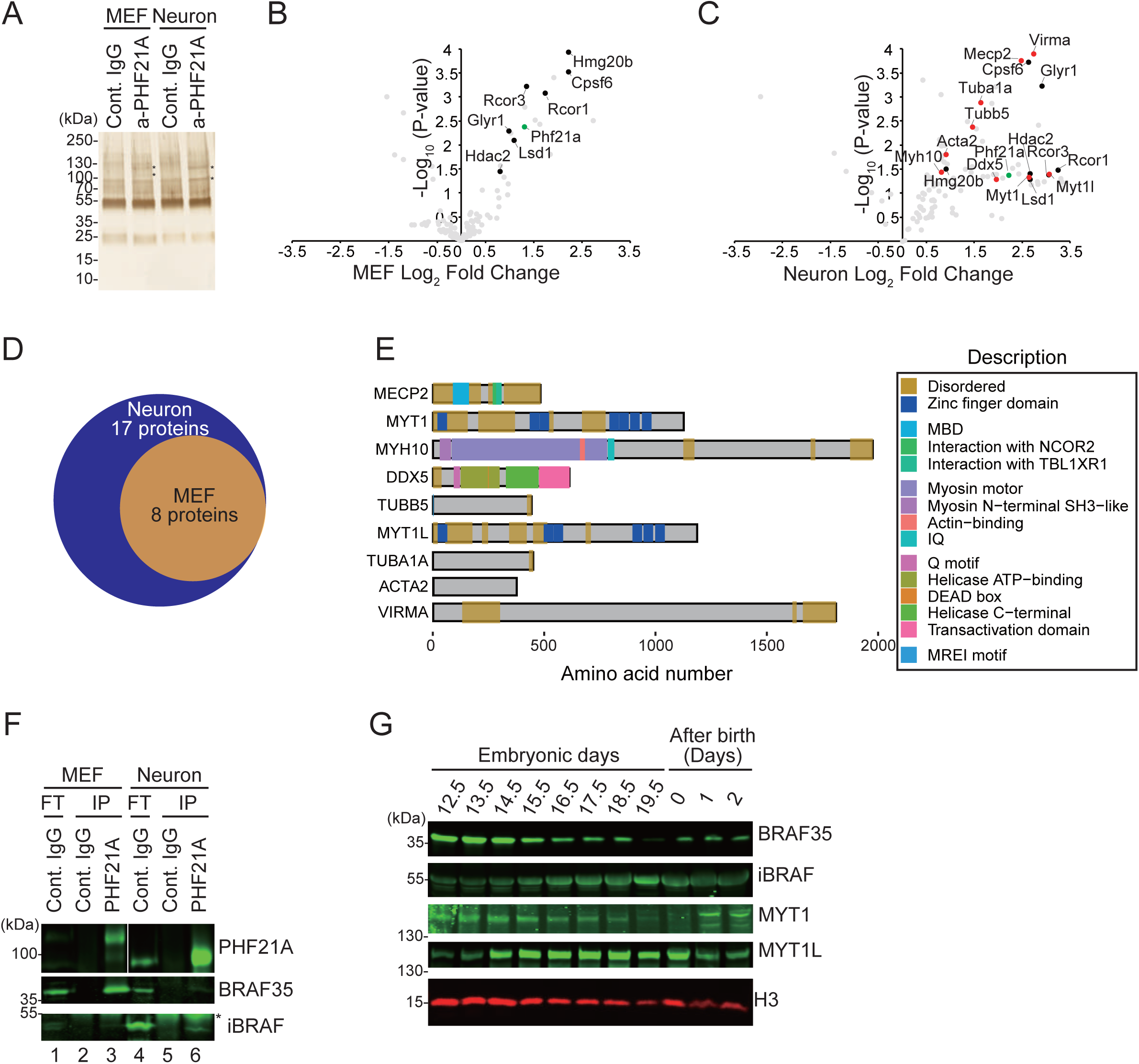
Proteomics analysis of PHF21A-containing complex in MEFs and neurons. **(A)** Silver staining of proteins co-precipitated by an anti-PHF21A antibody. Nuclear extracts from MEFs and cortical neurons (DIV7) were used. Asterisks indicate proteins specifically found in PHF21A-immunoprecipitates, **(B-C)** Co-IP-MS analysis of PHF21A-associated proteins in MEF **(B)** and cortical neurons **(C)**. Volcano plots of identified proteins (n=3). Green dots: the bait protein, PHF21A. Black and red dots: the interactors (Cutoff: FC > 1.5, *P*-value < 0.1, and peptide ≥ 6). Red dots: neuron-specific interactors with the same cut-off. Gray dots: proteins that did not pass the cutoff. The x-axis shows the log_2_ FC (a-PHF21A antibody/control IgG), and the y-axis denotes −log_10_ *P*-values. **(D)** Venn diagram of PHF21A interactor in MEFs (orange) versus neurons (blue). **(E)** Domain organization of neuron-specific PHF21A interactors. **(F)** Co-IP-Western assays to test the interaction between PHF21A, BRAF35, or iBRAF. Asterisks indicate nonspecific bands. **(G)** Expression of neuron-specific PHF21A interactors in the developing mouse brains. Whole-cell lysates prepared from mouse whole brains (E12.5 and E13.5) and cortices (from E14.5 to after birth 2) at indicated periods were subjected to Western blot analysis using antibodies as indicated.

**Figure 5:**
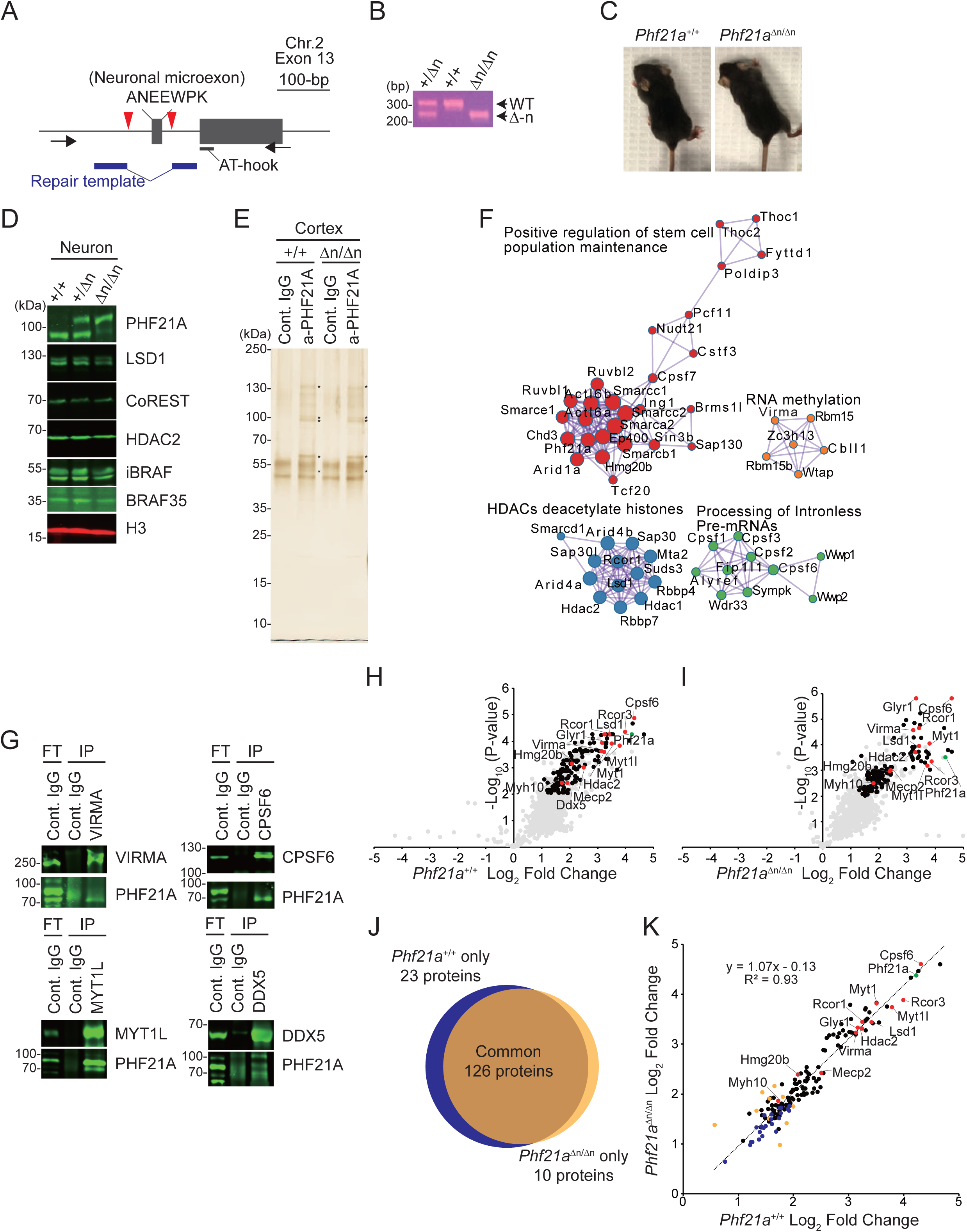
Binding partners of PHF21-n and ectopic PHF21A-c in neurons are highly similar. **(A)** Generation of *Phf21a*-neuronal exon knockout allele (Δn). Black arrows: PCR primers. Red triangles: CRISPR cut sites. **(B)** Genotyping PCR analysis of *Phf21a*^+/+^, *Phf21a*^+/Δn,^ and *Phf21a*^Δn/Δn^ mice using the primer sets shown in Fig. 5B. **(C)** Representative picture of P15 *Phf21a*^+/+^ (left) and *Phf21a*^Δn/Δn^ (right) mice. **(D)** Western blot analysis of lysates prepared from cortical neurons (DIV7) isolated from *Phf21a*^+/+^, *Phf21a*^+/Δn,^ and *Phf21a*^Δn/Δn^ mice using antibodies as indicated. PHF21A-c instead of PHF21A-n is expressed in *Phf21a-n*^Δn/Δn^ neuron. **(E)** Silver staining of proteins co-precipitated by anti-PHF21A antibody using the nuclear extracts from P0 cortices of the indicated genotypes. **(F)** Functional enrichment analysis of PHF21A-interacting proteins with Metascape (41). The statistically significant PHF21A interactors in *Phf21a*^+/+^ cortices were used. Log_10_ *P*-value < −10. The molecular networks that contain proteins validated by reciprocal IP experiment are presented. **(G)** Reciprocal Co-IP-Western assays to validate the interaction between PHF21A and newly-identified interaction partners. **(H and I)** Volcano plots of Co-IP-MS analysis of PHF21A-interacting proteins (n=3-4) using *Phf21a*^+/+^ **(H)** and *Phf21a*^Δn/Δn^ **(I)** cortices. Green dots: the bait protein, PHF21A. Black and red dots: statistically significant interactors (Cutoff: FC > 2, *Padj*-value < 0.01, and peptide ≥ 6). Red dots: neuron-specific proteins identified in Fig. 4C that pass the cutoff. Gray dots: proteins that do not pass the threshold. The x-axis shows the log_2_ FC (a-PHF21A antibody/control IgG), and the y-axis denotes −log_10_ *P*-values. **(J)** Venn diagram of PHF21A interactor in *Phf21a*^+/+^ cortex (Blue) versus *Phf21a*^Δn/Δn^ cortex (orange). **(K)** Scatter plot comparing the log2FC (a-PHF21A antibody/control IgG) of all PHF21A-associated proteins between WT and *Phf21a*^Δn/Δn^ cortices. The R^2^ value and the fit of the linear regression are indicated. Green dots: the bait protein, PHF21A. Black and red dots: statistically significant interactors in *Phf21a*^+/+^ a-PHF21A and *Phf21a*^Δn/Δn^ (Cutoff: *Padj*-value < 0.01, and peptide ≥ 6). Red dots: neuron-specific proteins identified in Fig. 4C that pass the cutoff. Blue dots: statistically significant interactors in *Phf21a*^+/+^ a-PHF21A. Orange dots: statistically significant interactors in *Phf21a*^Δn/Δn^. The x-axis shows the log_2_ FC of *Phf21a*^+/+^ (a-PHF21A antibody/control IgG). The y-axis shows the log_2_ FC of *Phf21a*^Δn/Δn^ (a-PHF21A antibody/control IgG).

BRAF35/ HMG20B is a component of the canonical PHF21A-LSD1 complex purified from non-neuronal HeLa cells (7, 8). Previous work showed that a BRAF35 paralogue called iBRAF (aka. HMG20A) is expressed in mature neurons instead of BRAF35 and inhibits the action of BRAF35, thereby promoting the expression of neuron-specific genes (36, 37). The dominant expression of BRAF35 in neuroprogenitors and iBRAF in mature neurons led us to postulate that BRAF35 interacts with PHF21A-c while iBRAF can interact with PHF21A-n. Our MS analysis identified BRAF35 in both MEF and neurons, yet iBRAF was not detected in neurons. We reasoned that iBRAF was not detected due to its small size. We, therefore, tested whether BRAF35 and/or iBRAF interact with PHF21A isoforms by Co-Immunoprecipitation Western analysis (Co-IP-WTN). We found that BRAF35 was expressed at a comparable level in MEF and neurons, whereas the iBRAF level was significantly higher in neurons compared to MEF (Fig. 4F, lanes 1&4). In the MEF PHF21A-IP samples, BRAF35 interacted with PHF21A-c as expected, while in the neuronal PHF21A-IP samples, iBRAF was readily detected, and a weak BRAF35 signal was detected. These results suggest that BRAF35 can interact with PHF21A-n; however, iBRAF competes with BRAF35 for PHF21A-n binding in neurons.

Having identified neuron-specific PHF21A-interacting proteins, we sought to understand the temporal dynamics of PHF21A-containing complexes during brain development. To this end, we examined the expression kinetics of these novel PHF21A partners in developing mouse brains by Western blot analyses. Consistent with the previous reports (36, 37), BRAF35 expression decreased during brain development, which is accompanied by the concomitant increase of iBRAF expression (Fig. 4F). Such reciprocal expression pattern was also found in MYT1 and MYT1L; the MYT1 level was higher in early development (E12.5 to E15.5), whereas MYT1L started increasing from E14.5. After birth, however, the expression of BRAF35 and MYT1 returned, while their pairs, iBRAF and MYT1L, plunged their expression again, representing an intricate regulation of these complex components. MeCP2 level increased after birth, consistent with its expression coinciding with synaptogenesis (38, 39). Together, these results suggest that PHF21A interacts with both canonical and additional neuron-specific proteins, and the interaction partners can change during neuronal differentiation owing to their dynamic expression patterns.

Unlike the previous report, we did not detect SVIL protein in the neuronal PHF21A complex, which might be the reason why we did not detect the H3K9 demethylation activity of the complex (Fig. 3C). Another work from our group demonstrates that neuronal splicing of PHF21A and LSD1 both contribute to the reduced H3K4 demethylation activity of the complex by interfering with their contacts with the nucleosomes (6). However, the reconstituted neuronal complex with LSD1-n, CoREST, and PHF21A-n still exhibited H3K4 demethylase activity, albeit weaker than its canonical counterpart (6). A plausible explanation of the difference is that the reconstituted complex in the other study was much more abundant than the complex from the brain used in the present study. Nonetheless, these observations all agree with the negative impact of the two neuronal splicing events on the enzymatic activity of the LSD1-PHF21A complex.

### The binding partners of PHF21A-n and ectopic PHF21A-c in neurons are highly similar

Having identified new neuron-specific PHF21A-binding partners, we wondered if being the neuronal form enables PHF21A to interact with these new partners in neurons. To address this question, we have generated a mouse model in which the *Phf21a* neuronal microexon is deleted by CRISPR-Cas9 (The *Phf21a*-Δn allele, Fig. 5A and 5B). In contrast to *Phf21a*-knockout (KO) mice, which lack both canonical and neuronal forms and die neonatally (40), homozygous *Phf21a*-Δn mice (*Phf21a*^Δn/Δn^) were viable (Fig. 5C). In the cortical neurons isolated in *Phf21a*^Δn/Δn^ mice, PHF21A-c is expressed at a comparable level with PHF21A-n in wildtype (*Phf21a*^+/+^) neurons (Fig. 5D). Thus, *Phf21a*-Δn represents the swapping mutant, in which PHF21A-n is replaced with PHF21A-c in neurons, allowing us to test if being neuronal form is necessary to interact with neuronal binding partners.

First, we examined whether the ectopic PHF21A-c expression in neurons alters the stability of binding partners for which specific antibodies are available. We did not observe any overt changes in the levels of LSD1, CoREST, HDAC2, BRAF35, and iBRAF, indicating that ectopic PHF21A-c does not impact the stability of known binding partners (Fig. 5D). In addition, overexpression of PHF21A-n, PHF21A-c, LSD1-n, or LSD1-c did not change the protein levels of endogenous PHF21A, LSD1, BRAF35, HDAC2, CoREST, and stably expressed iBRAF in 293T cells (Fig. S2A). Thus, neuronal PHF21A and LSD1 splicing events do not influence the stability of the binding partners examined.

Next, we performed proteomics quantification of PHF21A-associated proteins in *Phf21a*^+/+^ and *Phf21a*^Δn/Δn^ mice cortex (P0) with IP-MS (n=3 to 4 per genotype). As discussed earlier, at this stage of brain development, 94% of PHF21A and 81% of LSD1 are neuronal forms in *Phf21a*^+/+^ mice cortex (Fig. 1F and Fig. S1D). Thus, *Phf21a*^+/+^ data mostly represents PHF21A-n-associated factors in neurons, and *Phf21a*^Δn/Δn^ data represents PHF21A-c-associated factors in neurons. In this P0 brain proteomics study, unlike the earlier study with DIV7 cortical neuron cultures (Fig. 4A), we saw clear protein bands unique to PHF21A-IP samples (Fig. 5E) and identified many more proteins (149 in *Phf21a*^+/+^ samples, padj < 0.01, FC > 2, and peptide number ≥ 6) (Table S2). The greater numbers of identified proteins are likely due to higher PHF21A levels in the P0 brain compared to DIV7 neurons. With Metascape analysis (41), we found these PHF21A-interacting proteins belong to several protein-interaction networks: positive regulation of stem cell population maintenance, HDACs deacetylate histones, Processing of Intronless Pre-mRNAs, and RNA methylation (Fig. 5F and Table S3). Reciprocal IP assays validated some of these new PHF21A-interaction partners, including VIRMA, CPSF6, MYT1L, and DDX5 (Fig.5G).

The proteomic profile of the *Phf21a*^Δn/Δn^ mice cortex sample was highly similar to that of *Phf21a*^+/+^ (Fig. 5H and I). In the *Phf21a*^Δn/Δn^ mice cortex samples, we identified 136 proteins, which largely overlapped with those found in WT samples (Fig. 5J). 23 proteins passed the statistical threshold of interaction only in WT, and 10 proteins passed the threshold only in *Phf21a*^Δn/Δn^. To determine whether these differential binding factors between WT and *Phf21a*^Δn/Δn^ samples reflect genuine changes or borderline statistical significances, we compared the PHF21A-Ab / control IgG fold enrichment values (log2FC, the average of 3-4 replicates) of all PHF21A-interacting proteins found in either *Phf21a*^+/+^ or *Phf21a*^Δn/Δn^ samples (Fig. 5K). The log2FC enrichment values show a high concordance (R^2^ = 0.93) between genotypes, and the linear regression slope was 1.07. The log2FC values of these *Phf21a*^+/+^-and *Phf21a*^Δn/Δn^-specific binding proteins still show high concordance between genotypes, and these proteins mostly had low FC values. Furthermore, differential enrichment analysis between PHF21A-Ab-bound proteins between *Phf21a*^+/+^ and *Phf21a*^Δn/Δn^ yielded no statistically significant proteins (Fig. S3A). Thus, we concluded that interacting proteins did not substantially change in neurons when PHF21A-n was replaced with PHF21A-c.

These results demonstrate that PHF21A participates in unexpected molecular networks, such as post-transcriptional regulation, in neurons, and the PHF21A neuronal splicing is dispensable for the interaction between PHF21A and its neuronal partners.

## Discussion

In this work, we have described the asynchronous neuronal splicing of LSD1 and PHF21A in differentiating human neurons and developing mouse brains. The lagged LSD1 neuronal splicing implicated the step-wise transition of the complex abundance from LSD1-c + PHF21A-c, LSD-c + PHF21A-n, and then LSD1-n + PHF21A-n during brain development. The resulting complexes are deactivated for H3K4 demethylating activity along with this transition. Our proteomics studies revealed that PHF21A-n interacts with the known proteins and new neuron-specific partners in neurons. Unexpectedly, the neuronal splicing of PHF21A was dispensable for interacting with these neuron-specific binding partners.

The substrate specificity of LSD1-n has been shown as H3K4 (6, 15), H3K9 (16), and H4K20 (17). In the present study, similar to our prior study (6), we did not detect H3K9 or H3K20 demethylating activity in either the intermediate or mature PHF21A-LSD1 neuronal complexes. Instead, we observed the step-wise deactivation of the complex, which ultimately became incapable of reversing H3K4me in our assays. The lack of activity towards H3K9 can be explained by the absence of SVIL protein in our PHF21A-IP samples. We do not have any reasonable explanation for the lack of activity toward H4K20, given that this activity was observed in a purified system involving recombinant CoREST and LSD1 isoforms (17). It could be argued that H4K20 demethylation is not the function of LSD1-n that forms a complex with PHF21A; our complex was purified by anti-PHF21A antibody rather than anti-LSD1 antibody. However, our prior work did not detect H4K20 demethylation activity using recombinant LSD1-n and CoREST only either (6). It is imperative to identify the responsible variables between these observations and seek additional evidence, such as structural biology data, to resolve this issue fully.

Regardless of the LSD1-n neofunctionalization issue, the common conclusion agreed by all the studies is that LSD1 neuronal splicing impairs its H3K4 demethylation activity. In the present study, we further show that the deactivation of the H3K4-demethylating complex occurs in a stepwise fashion by two splicing events in LSD1-n and PHF21A-n during neurodevelopment. In addition, we also found that overall LSD1 and PHF21A levels decrease substantially from the late gestation period in the developing mouse brain (Fig. 1D & 1E). These observations indicate that the developing brain relentlessly decommissions the LSD1-PHF21A complex by multiple means: reducing its dose and intrinsic activity via mRNA splicing. The outstanding question to be addressed in the future is why neurons need to deactivate the complex. The swapping mouse mutant, *Phf21a*-Δn, will be a valuable resource to address this question.

However, our data also implicate that LSD1-PHF21A complex is not entirely eliminated from neurons (Fig.1 & Fig. 4). The remaining complex, containing LSD1-n and PHF21A-n, might play distinct functions by interacting with unique proteins in neurons. For example, we identified neuron-specific zinc-finger transcription factors MYT1 and MYT1L as neuron-specific PHF21A-binding proteins (Fig. 4). MYT1 and MYT1L belong to Myt-transcription factor family, which additionally involves MYT3 and expressed specifically in neurons with particularly high levels in the embryonic brain (42). Both MYT1 and MYT1L bind to a consensus DNA motif AAAGTTT (25) and transcriptionally suppress the Notch signaling pathway, which negatively regulates neurogenesis and, in turn, promotes neurogenesis (43, 44). MYT1L also represses deeper layer neuron transcriptional program in the adult prefrontal cortex (45) and non-neuronal genes (44, 46). SIN3-HDAC is a major corepressor that achieves MYT-mediated transcriptional suppression in vertebrates (45, 47). However, a MYT1 proteomics study with Neuro2A cell identified LSD1-CoREST-PHF21A complex (48), which is consistent with our results and suggests plausible roles of the complex in the MYT-mediated transcriptional regulation of neurodevelopmental gene expression program. Furthermore, MYT1L can also activate the transcription of neuronal maturation genes with unknown mechanisms (25, 49). It is tempting to speculate that the neuronal splicing of LSD1 and PHF21A is responsible for the functional switch of MYT1L from repressor to activator, with the neuronal LSD1-PHF21A complex acting as a dominant negative machinery. Lastly, this is the first report describing the interaction between the LSD1-PHF21A complex and post-transcriptional mRNA processing machinery such as CPSF6 and VIRMA. The association of these mRNA processing proteins is specific or much stronger in neurons (Figure 4 & 5). While CPSF6 controls alternative polyadenylation (50), VIRMA promotes m6A RNA methylation (51). Thus, neuronal LSD1-PHF21A complex, which largely lack H3K4 demethylation activity, might be involved in mRNA processing in neurons. Further investigations are needed to better understand this process.

## Experimental procedures

### Cell culture

#### LUHMES cells

LUHMES cells were purchased from ATCC (CRL-2927). The culturing and handling procedures of LUHMES cells were as described previously (52). In brief, LUHMES cells were grown at 37°C in a 5% CO_2_ atmosphere in DMEM/F-12 (Gibco 11330-032) containing N2 supplement (Gibco 17502048) and 40 ng/ml bFGF (Peprotech 100-18B). Before plating, plates were coated with PDL overnight at 37°C and washed with water thrice. Confluent cultures were passaged by trypsin digestion. One day after plating, the proliferation medium was replaced with DMEM/F-12 containing N2 supplement, 1 μg/ml tetracycline (Sigma T7660), 1 mM dibutyryl cyclic AMP (Selleck Chemicals S7858) and 2 ng/ml Recombinant Human GDNF (Peprotech 450-10).

#### HEK293T

HEK293T cells were grown at 37°C in a 5% CO_2_ atmosphere in DMEM (Gibco 11995065), supplemented with 10% FBS. **Primary Neurons**: The primary neuron culture was performed as previously described (6) with slight modifications. For molecular analyses, Cytarabine (10 *μ*M, aka AraC, Tocris) was added to the culture at DIV3 to eliminate the non-neuronal cell growth. **MEFs**: Mouse embryonic fibroblasts (MEFs) culture was performed as previously described (6). Cells were passaged less than twice for most experiments.

### Antibodies and PCR primers

Primary antibodies used are the following: rabbit anti-PHF21A (1:1000, in house, raised against 1-100 aa of human PHF21A expressed and purified from E.coli), rabbit anti-LSD1 (1:1000, Abcam ab17721), rabbit anti-BRAF35 (1:1000, ABclonal A4408), rabbit anti-HDAC2 (1:200, Santa Cruz sc-7899), rabbit anti-CoREST (1:1000, Abcam ab183711), mouse anti-H3 (1:500, Santa Cruz sc-517576), rabbit anti-H3K4me1 (1:5000, Abcam ab176877), rabbit anti-H3K4me2 (1:5000, Revmab bioscience 31-1037-00), rabbit anti-H3K9me1 (1:5000, Epicypher 13-0029), rabbit anti-H3K9me2 (1:5000, Abcam ab194680), rabbit anti-H4K20me1 (1:2500, Abcam ab9051), rabbit anti-H4K20me2 (1:5000, Abcam ab9052), mouse anti-H4 (1:200, Abcam ab31830), rabbit anti-iBRAF (1:1000, ABclonal A7286), rabbit anti-Myt1 (1:500, Invitrogen PA5-85510), rabbit anti-Myt1l (1:500, Proteintech 25234-1-AP) and rabbit anti-MeCP2 (1:500, Proteintech 10861-1-AP), Virma (1:1000, 25712-1-AP), CPSF6 (1:1000, ab99347), DDX5(1:2000, ab126730). The primers used for the amplification are as follows: hLSD1_F1: GCTGTGGTCAGCAAACAAG, hLSD1_R3: ATATTCCTTGCATAGGGCGGTC, hPHF21A_F1: GCAGTGACATACCTAAACAGC, hPHF21A_R1: CCAGGATGGTGTTCTTCATTTC, hTBP_1: GATCTTTGCAGTGACCCAGC, hTBP_2: CGCTGGAACTCGTCTCACTA, mLSD1_F3: CCCACTTTATGAAGCCAATGGAC, mLSD1_R3: CAACCGGTTAAATTCTTGTTCTACC, mPHF21A_F1: CAGTCACTTACCTTAACAGCAC, mPHF21A_R1: TGCTGCTCTTCATCTCCATAC, mTbp_qPCR_F: TTCAGAGGATGCTCTAGGGAAGA, mTbp_qPCR_R: CTGTGGAGTAAGTCCTGTGCC, Phf21a-n-F2: ACAGACGCCCAGCACCTTTAG, Phf21a-n-R2: GTAAGGGCTCCAAACCCCAG

### RNA purification and complete amplicon sequence

Total RNA was isolated using the RNeasy Plus Mini Kit (QIAGEN 74134) according to the manufacturer’s protocol. The RNA concentration was determined using a BioSpectrometer (Eppendorf). Total RNA (1 *μ*g) was used to generate the ProtoScript II First Strand cDNA Synthesis Kit (Biorad E6560S) following the manufacturer’s instructions. PCR reactions were performed with KOD Hot Start DNA Polymerase (Sigma 71086-3) using a Mastercycler X50a (Eppendorf). Complete amplicon sequencing was performed by the CCIB DNA Core Facility at Massachusetts General Hospital (Cambridge, MA). Illumina-compatible adapters with unique barcodes were ligated onto each sample during library construction. Libraries were pooled in equimolar concentrations for multiplexed sequencing on the Illumina MiSeq platform with 2×150 run parameters. Upon completion of the NGS run, data were analyzed, demultiplexed, and subsequently entered into an automated de novo assembly pipeline, UltraCycler v1.0 (Brian Seed and Huajun Wang, unpublished).

### Quantitative RT-PCR

The expression of different target genes was validated by quantitative PCR (qPCR) using the 7500 Real-Time PCR Systems (Applied Biosystems). The reactions were performed with the Power SYBR™ Green PCR Master Mix (Applied Biosystems 4367659) as recommended by the manufacturer. Real-time PCR was performed with a hot start step of 50 °C for 2 min and 95 °C for 10 min, followed by 30 cycles of 95 °C for 15 s, 60 °C for 1 min, and analyzed with 7500 software (Applied Biosystems). The PCR efficiency was calculated by linear regression between Ct values and concentrations of human LSD1-c and LSD1-n cDNA in pENTR plasmids. The resulting PCR efficiencies (*E*) were 73.5% for LSD1-c and 70.8% for LSD1-n, where 100% defines a doubling of DNA copy number per one PCR cycle. The correction value was calculated as [(*E*_LSD1-c_ +1)^30 / (*E*_LSD1-n_ +1)^30]. The LSD1-c: LSD1-n ratio obtained by the complete amplicon sequence described above was multiplied by the LSD1-n value with the correction value (1.62).

### Western blotting analysis (WTN)

Whole-cell lysates were prepared from mouse whole brains (E12.5 and E13.5) and cortices (E14.5 to P2). Cells were lysed in radioimmunoprecipitation assay (RIPA) buffer (50mM Tris-HCl pH7.5, 150mM 5M NaCl, 0.5% Sodium Deoxycholate monohydrate, 0.1% sodium dodecyl sulfate (SDS) and 1% TritonX-100) supplemented with Protease Inhibitor (Roche, 11873580001). Extracted proteins were boiled for 10 min with 2X Laemmeli buffer (100mM Tris-HCl pH6.8, 4% SDS, 0.2% Bromophenol blue, 20% Glycerol and 5% β-mercaptoethanol) at 100°C. Proteins were separated by SDS-polyacrylamide gel electrophoresis (PAGE) electroblotted onto a polyvinylidene difluoride (PVDF) membrane (Millipore, IPFL00010). PVDF membrane was masked with Intercept Blocking Buffer (LI-COR, 927-70001) for 2 hrs at 4°C, incubated with primary antibodies in Intercept Blocking Buffer overnight at 4°C, washed with PBST (137 mM NaCl, 2.7mM KCl, 11.9 mM phosphates and 0.1% Tween20) and incubated with secondary antibodies in Intercept Blocking Buffer for 1 h at RT. After the final washes, fluorescence signals were detected using Odyssey DLx Imager (LI-COR).

### Co-immunoprecipitation analysis (Co-IP)

Nuclei were enriched from the Dounce homogenized 293T cells, MEFs, neurons, and cortex using EZ Nuclei Lysis buffer (10 mM Tris at pH 7.4, 10 mM NaCl, 5 mM MgCl2, 0.5% NP-40, x1 Protease inhibitor cocktail). Nucleoproteins were extracted from nuclei with the same volume of IP Extraction Buffer (20mM HEPES at pH 7.9, 1.5mM MgCl_2_, 0.6M KCl, 0.2mM EDTA, 0.5mM DTT, 25% Glycerol, x1 Protease inhibitor cocktail) as EZ Nuclei Lysis buffer for 30 min at 4°C. For Co-IP, samples were bound to 5ug of crosslinked antibody (Rabbit IgG, a-PHF21A) each sample for 3 hrs at 4°C. Then, samples were subjected to SDS-PAGE and Western blot analyses. For Co-IP WTN, to reduce the background signals originating from IgG molecules used for IP, the PHF21A antibody was biotinylated by Pierce Antibody Biotinylation Kit (90407) following the manufacturer’s protocol and used as the primary WTN antibody. Likewise, for PHF21A IP, antibodies (Control rabbit IgG, a-PHF21A) were crosslinked as follows: first, the antibodies were reacted with Protein A/G beads (1:1 mixture) overnight at 4 °C, then crosslinked by 10*μ*M DMP (Thermo 21667) in 0.2 M sodium borate pH 9.0 for 1 h at RT. The reaction was quenched with 0.2 M Tris-HCl pH 8.0 at room temperature for 1.5 hrs, and the antibody-conjugated beads were washed with IP buffer and used for IP reaction.

### Demethylation assays

The PHF21A-containing complexes were immunoprecipitated with 50 ug of crosslinked antibody for 3 hrs at 4°C. One ug of recombinant designer demethylated nucleosomes (EpiCypher) was incubated with the PHF21A-containing complexes from 293T cells, LUHMES cells, or the mouse cortices for 3.5 hours at 37°C in histone demethylation buffer (50 mM Tris-HCl at pH 8.0, 50 mM KCl, 0.5% BSA, 5% glycerol, 0.5 mM DTT). The demethylation activity was measured by the appearance of monomethylation in WTN using specific methyl-histone antibodies listed above.

### Co-IP-MS

The immunoprecipitated protein was isolated from MEF cells and DIV7 neurons of E16.5 or P0 mouse cortices with the abovementioned in-house PHF21A antibody. Rabbit IgG from unimmunized rabbits (Jackson ImmunoResearch) was used as a control. pus (n = 3∼4). The PHF21A-complex was immunoprecipitated with 40 ug of a-PHF21A antibody as described above, eluted by 0.1% trifluoroacetic acid, and neutralized by 200 mM EPPS (4-(2-hydroxyethyl) piperazine-1-propane sulfonic acid) buffer. Then, samples were added to a 1:1 mixture of 2X Laemmeli buffer boiled for 10 min at 100°C. Proteins were separated on a 4-20% SDS-PAGE gel (Biorad 4561096, 5671095) and visualized with a Piece Silver Stain kit (Thermo 24612) according to the manufacturer’s protocol. The samples were multiplex with isobaric tandem mass tags (TMT) and analyzed with Orbitrap Fusion Tribrid Mass Spectrometer at Thermo Fisher Scientific Center for Multiplexed Proteomics at Harvard Medical School (Cambridge, MA). Gene ontology (GO) annotation analysis was performed using Metascape (41).

### *Phf21a* exon 13 knockout (*Phf21a-Δn*) Mouse

CRISPR/Cas9 technology was used to introduce double-strand breaks upstream and downstream of *Phf21a* exon 13. The mouse *Phf21a* gene (ENSMUSG00000058318) is located on the forward strand of Chromosome 2 from nucleotides 92,014,451 to 92,195,011 (GRCm39). Loss of exon 13 will cause an out-of-frame mRNA with multiple premature termination codons expected to trigger nonsense-mediated mRNA decay (53). The CRISPR algorithm (54, 55) was used to identify highly specific single guide RNAs (sgRNA) in introns 12 and 13: intron 12 sgRNA (C189C): 5’-AAGGTTAATACACAGGCCAG (PAM=AGG)-3’ (CFD score of 85). Intron 13 sgRNA (C189Y): 5’-AAAATGATCTTACATACCTT (PAM=TGG)-3’, (CFD score of 69). Phosphorothioate-modified sgRNA was synthesized by Synthego (56, 57). Each sgRNA (30 ng/ul) was complexed with enhanced specificity (eSP) Cas9 protein (50 ng/ul from Millipore-Sigma (58) and individually validated for causing chromosome breaks in mouse zygotes. The ribonucleoproteins (RNPs) were microinjected into fertilized mouse eggs. Eggs were placed in culture until they developed into blastocysts. DNA was extracted from individual blastocysts for analysis. PCR with primers spanning the predicted cut sites was used to generate amplicons for Sanger sequencing (59). To test sgRNAs C189C and C189Y, a 668-bp amplicon was produced with forward primer: 5’-GTCTGAACTGTTAGCAAAGAGACACAGAAA-3’; and reverse primer 5’-AGAGAGTACATGTCCCCAAGTTACTTAC-3’. Sequencing electropherograms of amplicons from individual blastocysts were evaluated to determine if small insertions/deletions caused by non-homologous end joining (NHEJ) repair of chromosome breaks were present (60). Using high-specificity sgRNA and enhanced specificity Cas9 protein dramatically reduces the likelihood of off-target hits in mice (61). The CRISPR reagents were microinjected into fertilized mouse eggs produced by mating superovulated C57BL/6J female mice (Jackson Laboratory stock no. 000664) with C57BL/6J male mice as described (62). CRISPR/Cas9 microinjection of zygotes produced founder mice. Generation zero founder (G0) pups were identified by Sanger sequencing of the PCR amplicons spanning the expected deletion. G0 founders were mated with wild-type C57BL/6J mice to obtain germline transmission of the *Phf21a* mutant allele. *Phf21a-Δn* mice were backcrossed into C57BL/6J for at least five generations to remove off-target mutations.

### Mice

We bred the animals in groups of one or two under specific pathogen-free conditions (*ad libitum* access to food and water, 12:12 light: dark cycle, light on at 06:00 am). Mice were tagged using ear punch and randomly assigned to each experiment. Timed-pregnant CD-1 mice were purchased from Charles River to obtain E16 embryos and P0 pups. All animal use followed NIH guidelines and approved by the University of Michigan Committee on Use and Care of Animals.

### Expression plasmids

PHF21A-c, PHF21A-n, LSD1-c, and LSD1-n cDNAs cloned into pENTR-D-TOPO, the Gateway Entry System (Invitrogen) entry vector, were previously described (6). Full-length coding sequences of BRAF35 and iBRAF were synthesized (Twist Bioscience) and cloned into pENTR-D-TOPO. To prepare expression plasmids, the cDNA fragments in pENTR-D-TOPO were transferred to the modified pHAGE plasmids (6) using LR recombination (Invitrogen 56484).

### Establishment of a stable cell line expressing iBRAF and transfection

pHAGE plasmid containing iBRAF was co-transfected with psPAX2 and pMD2.G into 293T cells as previously described (63). After 2 days of passaging, 1 ug/mL of puromycin was added to the medium and selected for >3 days. HEK293T cells were transfected with expression plasmids carrying PHF21A-c, PHF21A-n, LSD1-c, LSD1-n, BRAF35 or iBRAF cDNAs using Lipofectamine 3000 reagent (Invitrogen) for 24 h and harvested.

## Data availability

The row data supporting this study’s findings are available at k06988349 in University of Michigan Deep Blue data.

## Supporting information

This article contains supporting information.

## Acknowledgments

We thank Drs. Uhn-Soo Cho and Sojin An for providing plasmids and helpful discussions, Thermo Fisher Scientific Center for the multiplexed proteomics study at Harvard Medical School, the CCIB DNA Core Facility housed in the Center for Computational and Integrative Biology at Massachusetts General Hospital for the complete amplicon sequence service, and the members of the Iwase laboratory for their helpful discussions. We acknowledge Wanda Filipiak, Galina Gavrilina, and the Transgenic Animal Model Core of the University of Michigan’s Biomedical Research Core Facilities for of designing and producing the *Phf21a* mutant mice.

## Author contributions

M.N. and R.S.P. investigation; M.N. and S.I. conceptualization; M.N. and S.I. methodology; M.N. formal analysis; M.N. visualization; T.L.S and E.H. resources; M.N. and S.I. writing; M.N. and S.I. project administration; M.N. and S.I. funding acquisition; S.I. supervision.

## Funding and additional information

This work was supported by National Institutes of Health (NIH) grants (R01NS116008, R21NS12544, R21MH127485 to S.I.) and the University of Michigan Pandemic Research Recovery Program (U078150, to M.N.). The work in the Michigan Transgenic Core was supported by the National Cancer Institute of the National Institutes of Health under award number P30CA046592.

---

**Supporting Figure 1:**
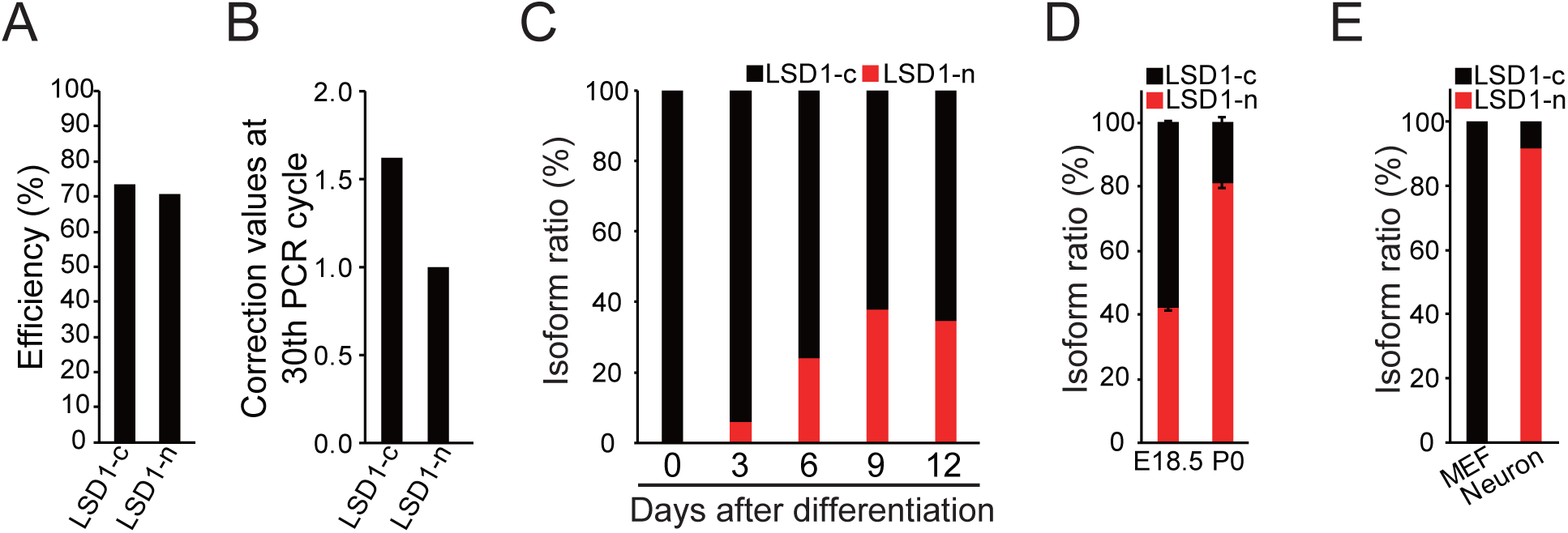
The difference in PCR efficiency in LSD1 isoform amplicons. **(A)** PCR efficiency of LSD1 mRNA isoforms empirically determined by the LSD1-c and LSD1-n cDNA-carrying plasmids. The values were averaged from three technical replicates at four concentrations. **(B)** Correction values of LSD1 isoform expression after 35 cycles using the efficiency difference determined in **(A)**. LSD1-c is estimated to have a 1.62-fold higher expression than RT-qPCR measurement after the correction. **(C)** The ratio of LSD1-c and LSD1-n mRNA isoforms. Cells were differentiated into neurons as indicated and harvested on day 3 to day 12. Data were adjusted for PCR efficiency determined in (B). **(D)** The ratio of LSD1-c and LSD1-n mRNA isoforms (n=3, mean ± S.E.M.) in the developing mouse brain. Data were adjusted for PCR efficiency determined in (B).

**Supporting Figure 2:**
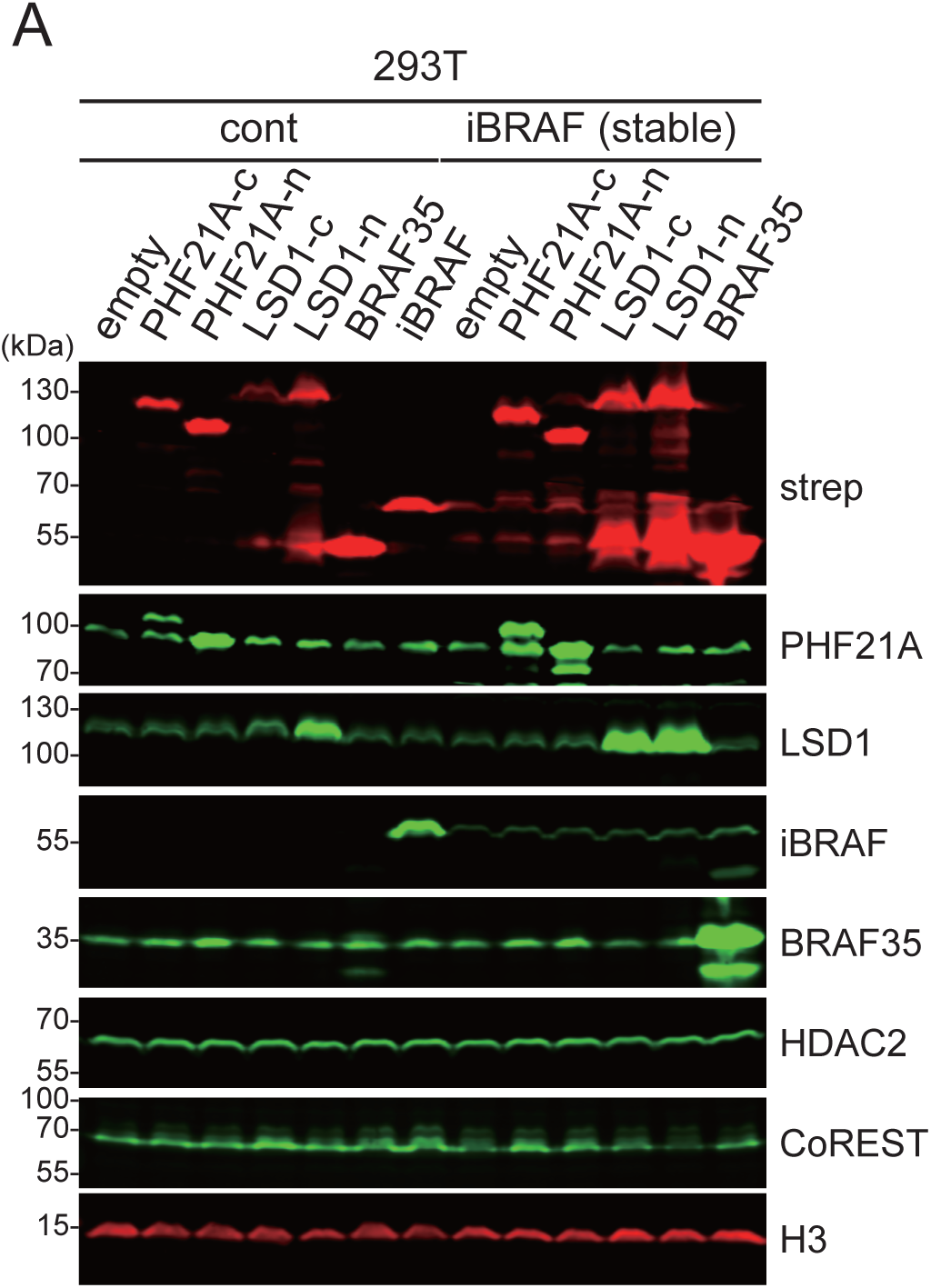
Impact of overexpressing PHF21A interactors on the stability of other complex components. **(A)** The protein levels of PHF21A-interactors, LSD1, iBRAF, BRAF35, HDAC2, and CoREST were examined after overexpressing the indicated proteins in 293T cells by Western blot analysis. 293T cells do not express iBRAF, so we generated a stable cell line expressing iBRAF and examined the impact of overexpression of the above molecules on the iBRAF protein level.

**Supporting Figure 3:**
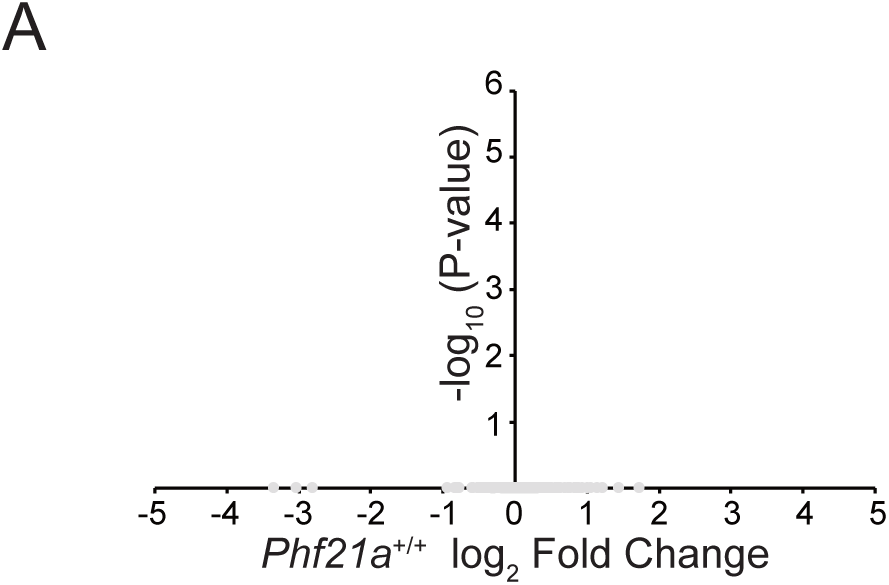
Binding partners of PHF21-n and ectopic PHF21A-c in neurons are highly similar. **(A)** Volcano plots of Co-IP-MS data comparing a-PHF21A IP samples between the *Phf21a*^+/+^ and *Phf21a*^Δn/Δn^ cortices (n=3-4). The x-axis shows the log_2_ FC (a-PHF21A IP of *Phf21a*^+/+^ cortices / a-PHF21A IP of *Phf21a*^Δn/Δn^ cortices), and the y-axis denotes −log_10_ *P*-values. Gray dots: proteins that do not pass the threshold.

**Supporting Table 1: Co-IP-MS analysis of PHF21A-associated proteins in MEF and cortical neurons.**

The complete list of identified proteins in Co-IP-MS comparing MEF and cortical neurons. Raw MS values (summed S/N values scaled to 100 per protein) and statistics, including −log(P), and log2FC are included.

**Supporting Table 2: Co-IP-MS analysis of PHF21A-associated proteins in *Phf21a*^+/+^ and *Phf21a*^Δn/Δn^ cortices.**

The complete list of identified proteins in Co-IP-MS comparing *Phf21a*^+/+^ and *Phf21a*^Δn/Δn^ cortices. Raw MS (summed S/N values scaled to 100 per protein) and values and statistics, including −log(Padj), and log2FC are included.

**Supporting Table 3: Functional enrichment analysis of PHF21A-interacting proteins with Metascape.**

Complete list of molecular networks PHF21A-interacting proteins participate. The analysis was done with Metascape.

